# AllergenAI: a deep learning model predicting allergenicity based on protein sequence

**DOI:** 10.1101/2024.06.22.600179

**Authors:** Chengyuan Yang, Surendra S Negi, Catherine H Schein, Werner Braun, Pora Kim

**Author notes:** contributed equally (Chengyuan Yang, Surendra S Negi). Address correspondence to the contact corresponding author: Pora Kim, Ph.D. School of Biomedical Informatics The University of Texas Health Science Center at Houston 7000 Fannin St., Houston, TX 77030, Werner Braun, Ph.D. Department of Biochemistry and Molecular Biology Sealy Center for Structural Biology and Molecular Biophysics University of Texas Medical Branch 301 University Blvd, Galveston, TX 77555-0304.

## Abstract

Innovations in protein engineering can help redesign allergenic proteins to reduce adverse reactions in sensitive individuals. To accomplish this aim, a better knowledge of the molecular properties of allergenic proteins and the molecular features that make a protein allergenic is needed. We present a novel AI-based tool, AllergenAI, to quantify the allergenic potential of a given protein. Our approach is solely based on protein sequences, differentiating it from previous tools that use some knowledge of the allergens’ physicochemical and other properties in addition to sequence homology. We used the collected data on protein sequences of allergenic proteins as archived in the three well-established databases, SDAP 2.0, COMPARE, and AlgPred 2, to train a convolutional neural network and assessed its prediction performance by cross-validation. We then used Allergen AI to find novel potential proteins of the cupin family in date palm, spinach, maize, and red clover plants with a high allergenicity score that might have an adverse allergenic effect on sensitive individuals. By analyzing the feature importance scores (FIS) of vicilins, we identified a proline-alanine-rich (P-A) motif in the top 50% of FIS regions that overlapped with known IgE epitope regions of vicilin allergens. Furthermore, using∼ 1600 allergen structures in our SDAP database, we showed the potential to incorporate 3D information in a CNN model. Future, incorporating 3D information in training data should enhance the accuracy. AllergenAI is a novel foundation for identifying the critical features that distinguish allergenic proteins.

## INTRODUCTION

Proteins from different plants, fungi, and animals can sensitize individuals and stimulate IgE-mediated allergic responses [1]. Symptoms range from rashes to difficulty breathing to potentially deadly anaphylaxis [2], as has been observed for peanut allergies [3, 4]. While desensitization by immunotherapy has made great strides, it is not an effective treatment for all individuals due to the need for repeated administration over months or years and the associated danger of anaphylaxis [5] [6]. This leaves many allergy sufferers facing avoidance of known triggers as the sole therapy. However, elimination even of known allergens is difficult. Similar proteins lurk in many seemingly unrelated sources, and there is always the danger of cross-reactions.

To avoid introducing new allergens into our environment, there is an urgent need for a deep learning model to define specific molecular features that render a protein allergenic [7]. We and others previously established that known allergenic proteins populate a small subfraction of protein families (Pfam) [8, 9]. However, not all proteins in these Pfams are allergenic [10]. Sequence identity alone is not adequate to explain cross-reactivity; e.g., while IgE binding proteins of peanut and tree nuts have a very low identity to each other, over 30% of individuals with a peanut allergy react strongly to walnuts [11–13]. We hypothesize that introducing a deep learning approach into the allergenic protein classification problem will give a deeper understanding of the allergenic protein features.

Here, we present a novel deep learning method, AllergenAI, to distinguish the allergenic potential of proteins (Figure 1). We used the collected data on protein sequences of allergenic proteins archived in three well-established resources, our SDAP 2.0 [14, 15], COMPARE [16] and AlgPred2 [17] to train a convolutional neural network (CNN) [18]. The prediction accuracy of the AllergenAI model was then assessed in a five-fold cross-validation with high accuracies in the training set and test data set. The prediction quality of AllergenAI is on par or superior to other machine learning methods, such as AllergenFP and AlgPred2. We also present the results of a pilot study to examine the extent of improvements which can be expected from including 3D structural information of proteins in the model. For that study we used the additional structural information of allergenic proteins in SDAP 2.0 in a CNN model and showed a significant, albeit small, improvement in the prediction qualities over using only amino acid sequences. We also suggest that future applications of AllergenAI can detect novel potential allergens in the proteomes of plants and animals.

**Figure 1.**
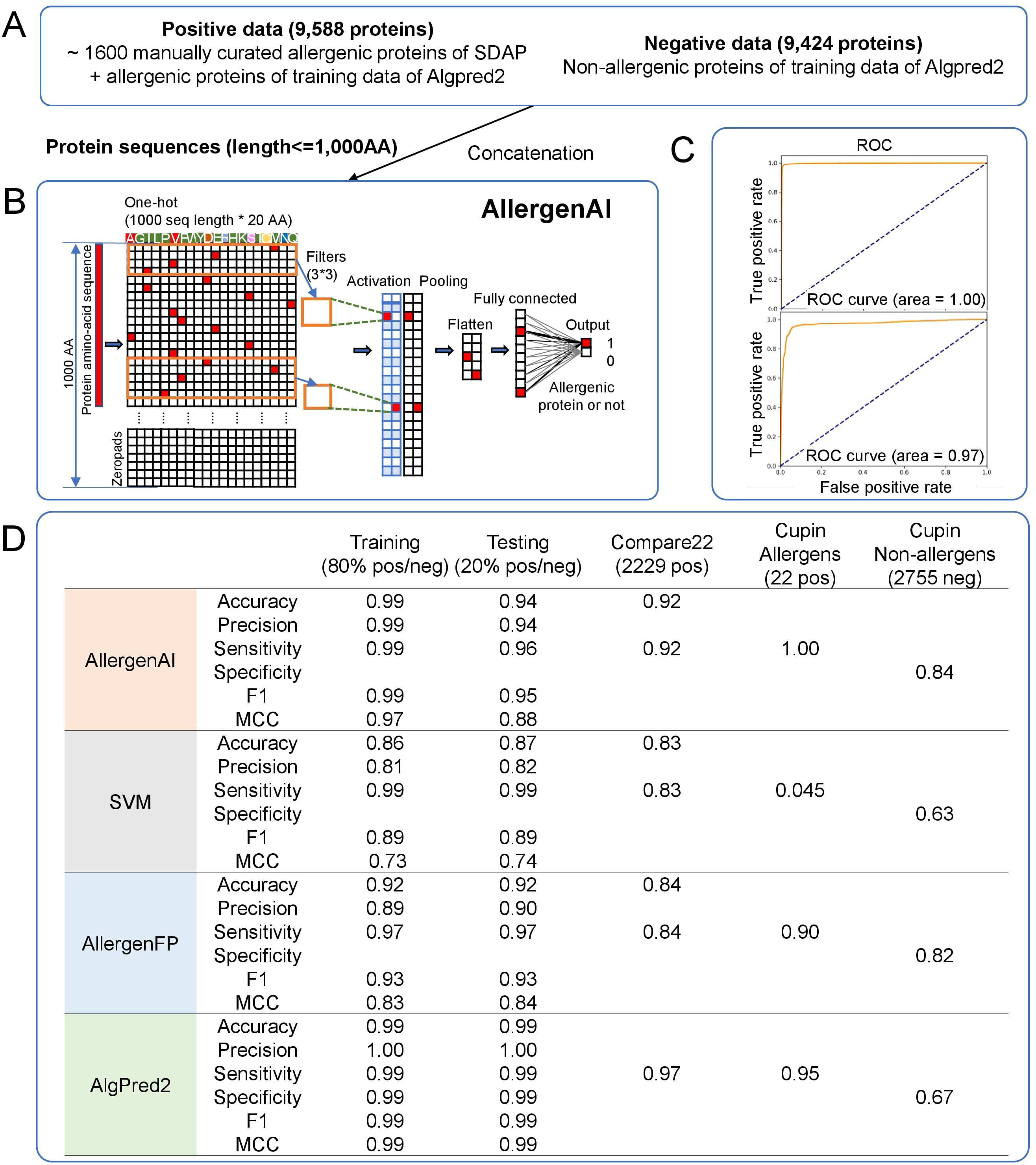
Overview and performance of AllergenAI. (A) Training and test data composition used to generate the CNN model of AllergenAI and evaluate its performance. (B) Input format and overview of the CNN model. (C) Receiver operating characteristic (ROC) curve for training and testing data sets. (D) Performance comparison with other machine learning approaches.

We validated Allergen AI by application to vicilins (also called 7S globulins), which are members of the very large, highly diverse Cupin family (PFAM) [19]. Several important food allergens in peanut, walnut, soybean and sesame seeds are vicilins, but there are also many non-allergenic homologues. The Cupin superfamily is characterized by a β-barrel 3D structure in their trimeric domains. Allergen AI detected four vicilin proteins as having allergenic potential from date palm, spinach, maize and red clover, which can be further assessed for IgE binding. This novel deep learning method could enrich current allergen databases with new information on allergenic sources that might affect sensitive individuals.

## RESULTS

### Overview of AllergenAI

Positive and negative datasets of allergens were collected from several, well-established databases that are available online. These data sources include our recently released, updated Structural Database of Allergenic Proteins (SDAP 2.0) with high quality 3D models predicted by AlphaFold2. SDAP 2.0 provided 1657 manually curated sequences including all allergens from the database of allergen nomenclature of the World Health Organization and International Union of Immunological Societies (WHO/IUIS), 334 experimentally determined X-ray or NMR structures, and 1565 3-D models. These are true allergens, integrated by manual curation. However, the small number of proteins might limit the capability of the CNN model to learn allergenic features from protein sequences. Therefore, we added allergenic protein sequence information available in various databases and online resources. AlgPred2 made their training data from COMPARE 2018, AllergenOnline, AlgPred, and reviewed SWISS-Prot allergens. We combined the training data sets of SDAP 2.0 and AlgPred2, for a total of 9,588 allergenic and 9,424 non-allergenic proteins for training (Figure 1A). The optimized weights generated by the deep learning process including activation, pooling, flattening, and fully connected functions resulted in a probability for each sequence to be allergenic (Figure 1B). The dimension reduction images show the better steadiness of the training data when we combine SDAP2.0 and AlgPred2 (Supplementary Figure S1). For our model, we randomly chose 80% for training and 20% for testing out of ∼ 9K positive and ∼ 9K negative datasets. Details on the training and test data sets are given in the Material and Methods section. For the parameters of AllergenAI, we set the Adam learning rate as 0.01, the batch size as 128, the number of epochs as 24, and used L2 regularization. The detailed model architecture is shown in Supplementary Figure S2. For the training set, the number of TP is 7542, the number of TN is 5906, the number of FP is 73, and the number of FN is 110. The accuracy of training prediction outputs is 0.99. The Precision and the Recall are 0.99, indicating that the AllergenAI correctly identifies 99% of known allergenic proteins (i,e. proteins belonging to the training set of allergenic proteins). For the testing set, the accuracy is 0.94, the precision is 0.94, and the recall is 0.96. The area under the curve (AUROC) was 1.00 and 0.97 for the training and testing datasets, respectively (Figure 1C).

### Performance comparison of AllergenAI to other machine learning models

We compared the performance of AllergenAI with three external validation datasets, which are allergens in the COMPARE 2022 dataset, vicilin allergens in the SDAP 2.0 database and non-allergens in the cupin family proteins. We first compared the performance of AllergenAI with another widely used machine learning model for binary prediction: SVM. We used the default parameters for the SVM model from the scikit-learn package. In general, AllergenAI has a better performance on all metrics for all datasets. The details of the numbers are shown in Figure 1D. Secondly, we compared AllergenAI with AllergenFP, a well-known model for predicting allergenicity. For the COMPARE 2022 dataset, which contains 2229 allergens, AllergenAI identified 2043 proteins correctly as allergenic (0.92) compared to AllergenFP with 0.84. For the cupin vicilin data, AllergenAI successfully identified all positive proteins and 2333 negative proteins which outperforms AllergenFP on these proteins. For the application to the known allergenic cupin sequences in SDAP and non-allergen sequences in CUPIN PFAM domain, AllergenAI outperformed also AlgPred2. Only AllergenAI is a primary protein sequence-based model while the other tools use curated allergenic features (i.e., sequence homology and physicochemical properties). In this aspect, we expected our approach may identify allergenic sequence features. All accuracy metrics are in Figure 1D. Figure 2 shows the Venn diagram of the comparison of the true negatively predicted proteins in the non-allergen sequence in cupin pfam domain. Among the three tools, AllergenAI had the highest true negative number (2332) among all three tools and performed well also in binary comparisons.

**Figure 2.**
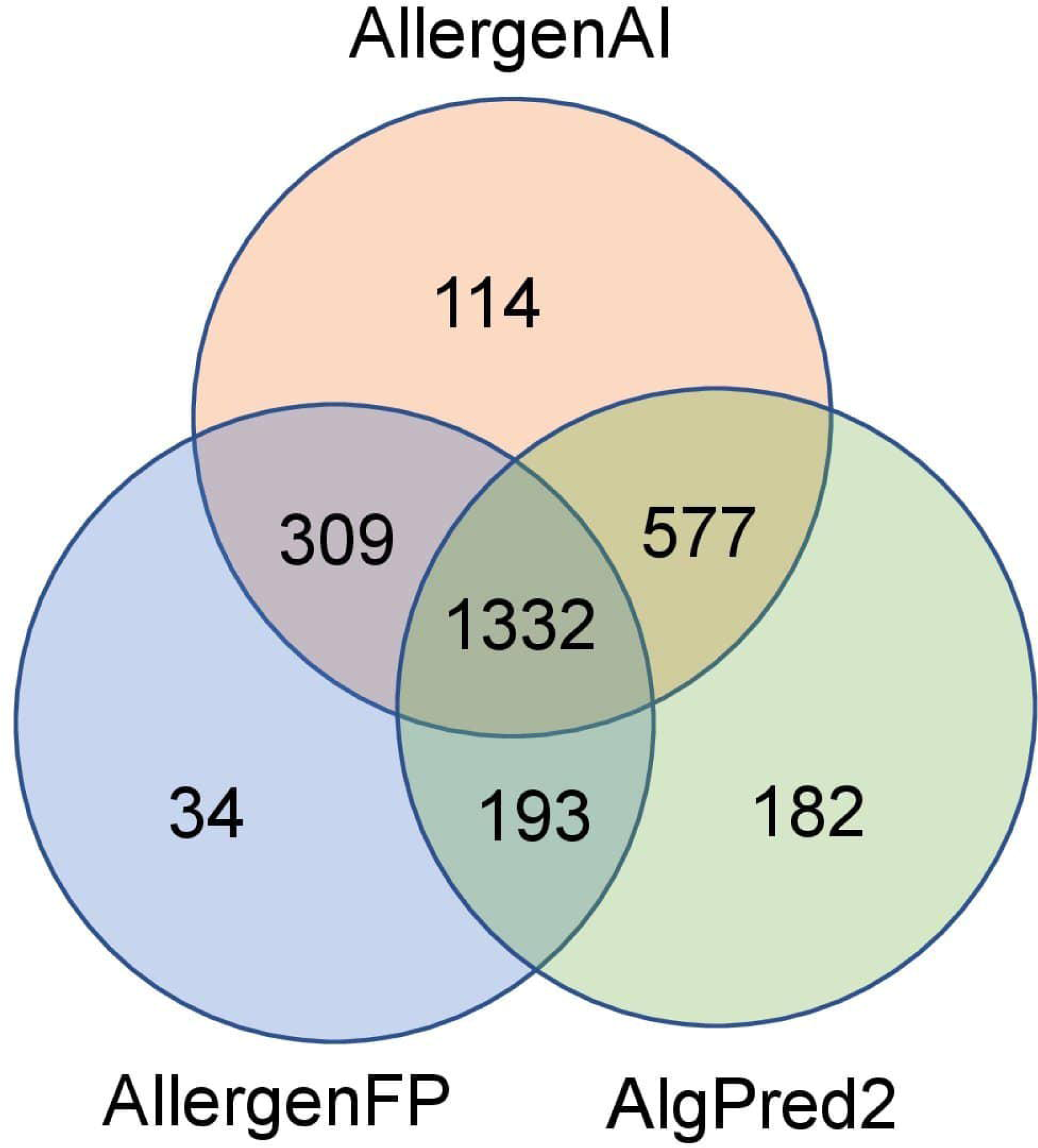
True negative prediction comparison among three tools in the non-allergens in cupin pfam proteins.

### Application to the proteins of vicilin family and feature importance score analysis identified proline-alanine–rich consensus sequence motifs

Allergens in the vicilin family are given in Table 2. Most are food allergens, seed storage proteins from tree nuts, sesame seeds or peanuts [19, 20]. The comparison of the predictions by the three tools, AllergenAI, AllergenFP and AlgPred2 to the individual allergens are given in Table 3. AllergenAI correctly predicted all of these vicilins as allergens, while AllergenFP missed two allergens, Mac I 1 andPin k 2 , and Algpred2 failed to correctly assign Coc n 1 with the default cutoff score of 0.3 (the programs default value). We next analyzed the feature importance scores (FIS) across the sequences of these vicilin allergens. For every 5 amino acids across protein sequence length, we masked as empty sequence and checked the different value of the AllergenAI output values (Figure 3A). Supplementary Figure S3 shows the histogram of the FIS of individual vicilin proteins. Figure 3B shows the distribution of the feature importance scores across individual vicilin protein sequence lengths. We analyzed all sequences which scored more than 50% feature importance using MEME [21]. MEME discovered novel, ungapped motifs (recurring, fixed-length patterns) in the top 50% of the feature importance scores regions detected by the scanning with AllergenAI. The most prominent of these was a proline-alanine (P-A) rich motif (Figure 4). A similar P-A motif was found to be the primary IgE binding epitope of a “long splice variant” protein implicated in primary chicken meat allergy [22]. A major IgE epitope in the propeptide of the house dust mite allergen Der p 3, NPILPASPNAT, called by the authors a proline-rich motif (PRM),unusual for a trypsin-like protease, is also P-A rich [23]. The importance of hydroxylated proline for IgE recognition was noted for another family of peanut allergens[24]. Thus the P-A rich feature recognized by Allergen AI is found in many major IgE epitopes.

**Figure 3.**
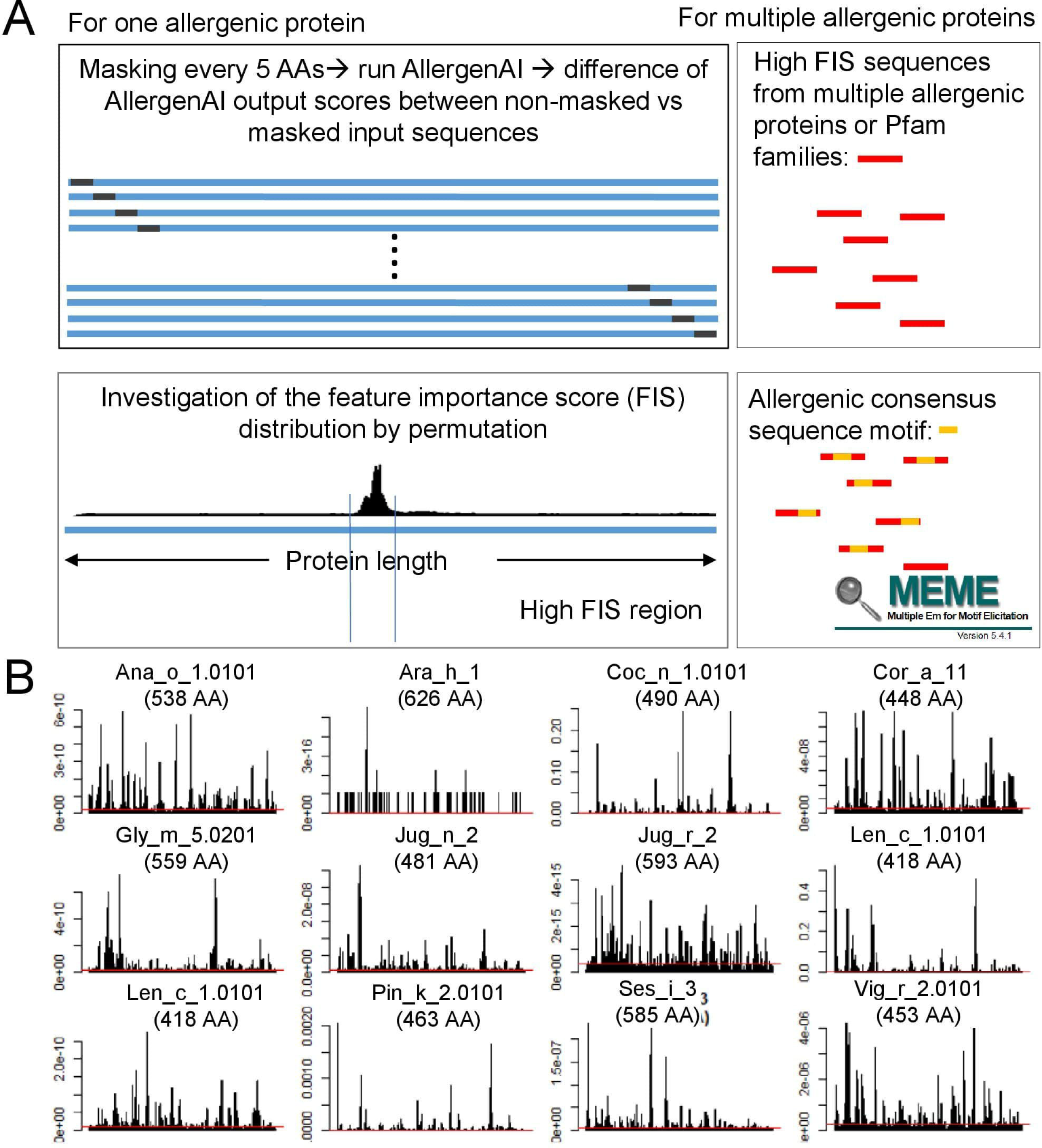
Analysis of the feature importance scores across the protein sequence lengths. (A) Conceptual explanation of our feature importance analysis. (B) Distribution of the feature importance scores across individual vicilin allergen protein sequence lengths.

**Figure 4.**
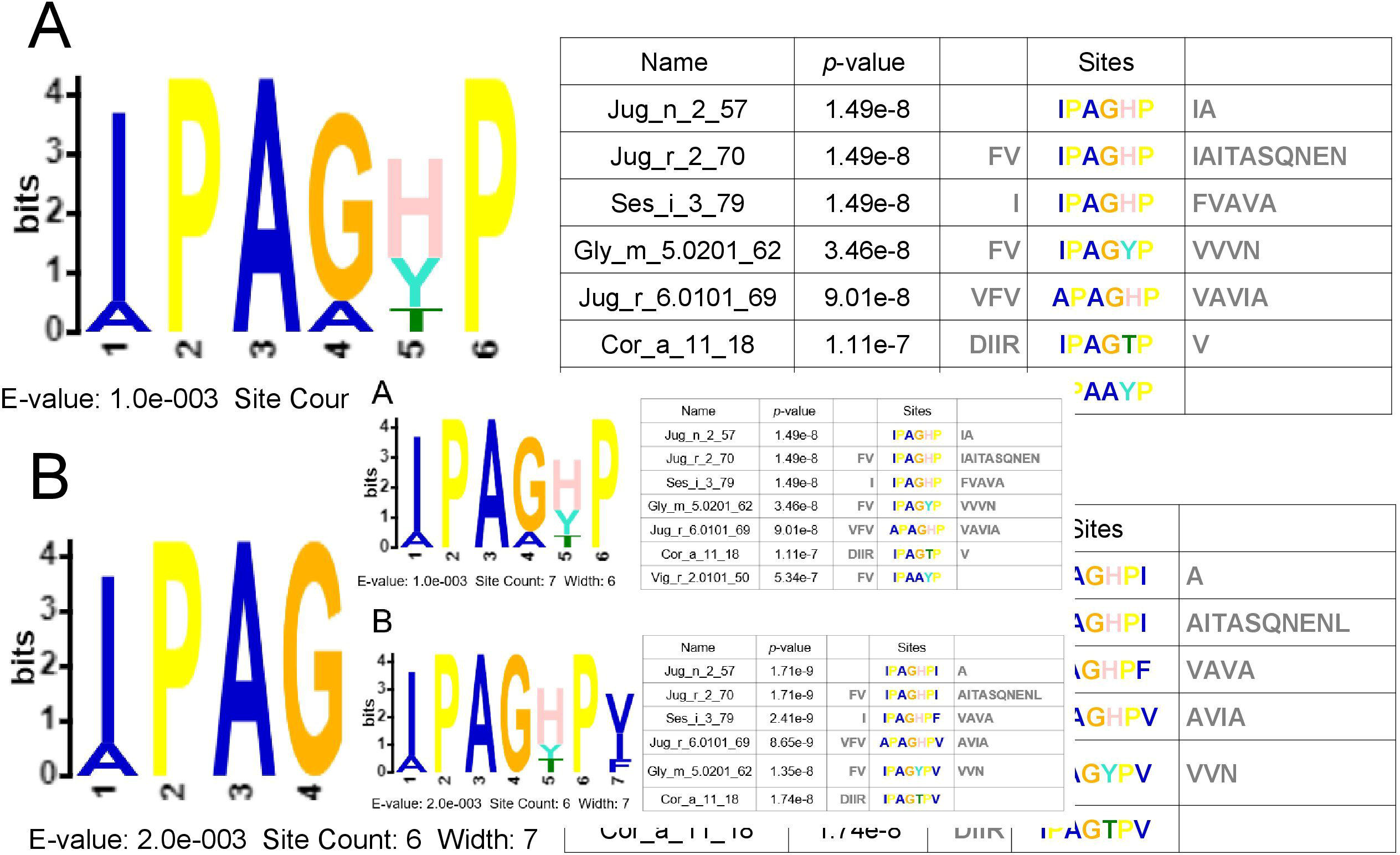
Consensus motif sequences in the top 50% FIS regions in vicilin proteins. We used MEME to identify the consensus protein sequence motif. (A) Significantly identified motif using min window size as 6 AA. (B) Significantly identified motif using min window size as 3 AA.

**Table 1.** Protein sequence distribution (# allergenic proteins) of all training/validation/ application data used in our study.

| AA length | # proteins per AA length |  |  |  |
| --- | --- | --- | --- | --- |
|  | All SDAP positive | PFAM | COMPARE | AlgPred2 |
| # all proteins | 1570 | 773 | 2237 | 2015 |
| <2000 | 1570 | 773 | 2237 | 2012 |
| <1900 | 1568 | 773 | 2237 | 2012 |
| <1800 | 1566 | 773 | 2237 | 2012 |
| <1700 | 1564 | 773 | 2235 | 2012 |
| <1600 | 1562 | 773 | 2233 | 2012 |
| <1500 | 1561 | 773 | 2233 | 2012 |
| <1400 | 1557 | 773 | 2233 | 2012 |
| <1300 | 1554 | 772 | 2230 | 2012 |
| <1200 | 1554 | 772 | 2230 | 2012 |
| <1100 | 1554 | 772 | 2230 | 2012 |
| <1000 | 1552 | 771 | 2229 | 2012 |
| <900 | 1552 | 771 | 2229 | 2011 |
| <800 | 1540 | 771 | 2213 | 2011 |
| <700 | 1526 | 771 | 2194 | 2011 |
| <600 | 1493 | 771 | 2154 | 2011 |
| <500 | 1433 | 738 | 2044 | 2011 |
| <400 | 1352 | 720 | 1928 | 1891 |
| <300 | 1171 | 658 | 1682 | 1553 |
| <200 | 871 | 620 | 1238 | 889 |
| <100 | 145 | 93 | 251 | 177 |
| pct at 400 AA | 0.861 | 0.931 | 0.862 | 0.938 |
| pct at 500 AA | 0.913 | 0.955 | 0.914 | 0.998 |
| pct at 600 AA | 0.951 | 0.997 | 0.963 | 0.998 |
| pct at 700 AA | 0.972 | 0.997 | 0.981 | 0.998 |
| pct at 800 AA | 0.972 | 0.997 | 0.981 | 0.998 |
| pct at 900 AA | 0.989 | 0.997 | 0.996 | 0.998 |
| pct a 1000 AA | 0.989 | 0.997 | 0.996 | 0.999 |
| max AA | 1915 | 1384 | 1773 | 3411 |

**Table 2 :** Allergens from the vicilin family in the SDAP database.

| Allergen Name | Order | Species | Route of Exposure | Common Name | Biochemical Name |
| --- | --- | --- | --- | --- | --- |
| Ana o 1 | Sapindales | Anacardium occidentale | Food | Cashew | Vicilin-like protein |
| Ara h 1 | Fabales | Arachis hypogaea | Food | Peanut | Cupin (Vicillin-type, 7S globulin) |
| Car i 2 | Fagales | Carya illinoensis | Food | Pecan | 2S albumin seed storage protein |
| Coc n 1 | Arecales | Cocos nucifera | Airway | Coconut | vicilin-like protein |
| Cor a 11 | Fagales | Corylus avellana | Food | Hazelnut | 7S seed storage globulin (vicilin-like) |
| Gly m 5 | Fabales | Glycine max | Food | Soybean | Beta-conglycinin (vicilin, 7S globulin) |
| Jug n 2 | Fagales | Juglans nigra | Food | Black walnut | Vicilin seed storage protein; 7S globulin |
| Jug r 2 | Fagales | Juglans regia | Food | English walnut | Vicilin (7S globulin) seed storage protein |
| Jug r 6 | Fagales | Juglans regia | Food | English walnut | 7S globulin seed storage protein, vicilin-like protein |
| Len c 1 | Fabales | Lens culinaris | Food | Lentil | Gamma-vicilin subunit |
| Lup an 1 | Fabales | Lupinus angustifolius | Food | Narrow-leaved blue lupin | Conglutin beta (7S seed storage globulin, vicilin) |
| Mac i 1 | Proteales | Macadamia integrifolia | Food | Macadamia | Vicilin |
| Pin k 2 | Pinales | Pinus koraiensis | Food | Korean pine | Vicilin |
| Ses i 3 | Lamiales | Sesamum indicum | Food | Sesame | 7S vicilin-like globulin |
| Vig r 2 | Fabales | Vigna radiata | Food | Mung bean | 8S Globulin (Vicilin) |
| Pis v 3 | Sapindales | Pistacia vera | Food | Pistachio | vicillin |

**Table 3.** Allergenicity prediction of vicilin proteins using three different methods.

| Protein name | AllergenAI |  | AllergenFP |  | AlgPred2 |  |
| --- | --- | --- | --- | --- | --- | --- |
|  | Output | Prediction | Output | Prediction | Output | Prediction |
| Ana_o_1.0101 | 1.00 | Allergen | 100 | Allergen | 1.00 | Allergen |
| Ana_o_1.0102 | 1.00 | Allergen | 100 | Allergen | 1.00 | Allergen |
| Ara_h_1 | 1.00 | Allergen | 100 | Allergen | 1.00 | Allergen |
| Ara_h_1 | 1.00 | Allergen | 100 | Allergen | 0.99 | Allergen |
| Car_i_2.0101 | 1.00 | Allergen | 100 | Allergen | 0.86 | Allergen |
| Coc_n_1.0101 | 1.00 | Allergen | 100 | Allergen | 0.02 | Non-Allergen |
| Cor_a_11 | 1.00 | Allergen | 100 | Allergen | 0.98 | Allergen |
| Fag_e_3.0101 | 1.00 | Allergen | 100 | Allergen | 0.99 | Allergen |
| Gly_m_5.0101 | 1.00 | Allergen | 100 | Allergen | 0.99 | Allergen |
| Gly_m_5.0201 | 1.00 | Allergen | 100 | Allergen | 0.99 | Allergen |
| Jug_n_2 | 1.00 | Allergen | 100 | Allergen | 0.98 | Allergen |
| Jug_r_2 | 1.00 | Allergen | 100 | Allergen | 1.00 | Allergen |
| Jug_r_6.0101 | 0.99 | Allergen | 100 | Allergen | 0.85 | Allergen |
| Len_c_1.0101 | 1.00 | Allergen | 100 | Allergen | 0.99 | Allergen |
| Len_c_1.0102 | 1.00 | Allergen | 100 | Allergen | 0.99 | Allergen |
| Lup_an_1.0101 | 0.99 | Allergen | 100 | Allergen | 1.00 | Allergen |
| Mac_i_1.0101 | 1.00 | Allergen | 100 | Allergen | 0.71 | Allergen |
| Mac_i_1.0201 | 0.99 | Allergen | -100 | Non-Allergen | 0.70 | Allergen |
| Pin_k_2.0101 | 1.00 | Allergen | -100 | Non-Allergen | 0.72 | Allergen |
| Ses_i_3 | 1.00 | Allergen | 100 | Allergen | 1.00 | Allergen |
| Vig_r_2.0101 | 1.00 | Allergen | 100 | Allergen | 0.96 | Allergen |
| Vig_r_2.0201 | 0.99 | Allergen | 100 | Allergen | 0.92 | Allergen |

### The allergenicity scores of AllergenAI correlate with real world data

We and others have emphasized that the number of protein families containing allergens is relatively small, compared to the variety of protein families in the protein universe [8, 9]. Literature surveys on food allergens in plants have indicated that physicochemical properties of allergens, such as thermal stability, resistance to proteolysis, or posttranslational modifications, are found prominently in allergens [25]. However, these properties are not universally true for all allergens and are difficult to use as criteria to decide if a new protein should be classified as an allergen. Also, several allergens, such as tropomyosins of shrimp and other crustaceans, can cause anaphylactic shocks in sensitized individuals but have high sequence identities to mammalian homologs that are not allergens.

We thus used our deep learning techniques to extract crucial features of allergens from the vast information in allergenic databases. We observed that a major basis of the prediction is the presence of small segments (three to four amino acids) of amino acids in these sequences that are also present in the allergens available in the SDAP database with moderate E-values. To test the significance of these small groups of amino acids, we calculated the feature importance scores (FIS) to distinguish proteins with allergenic potential. We previously showed that physico-chemical property (PCP) motifs based on PCP descriptors of the 20 amino acids, could distinguish allergens from random proteins[26]. The similarity of the segments with high FIS scores in the vicilin Ana o 3 allergen compared well to the location of PCP motifs[8] calculated from a sequence alignment of the vicilin family (Figure 5). This observation suggests that sudden changes in the FIS score identify segments in the protein sequences that are important for allergenicity.

**Figure 5:**
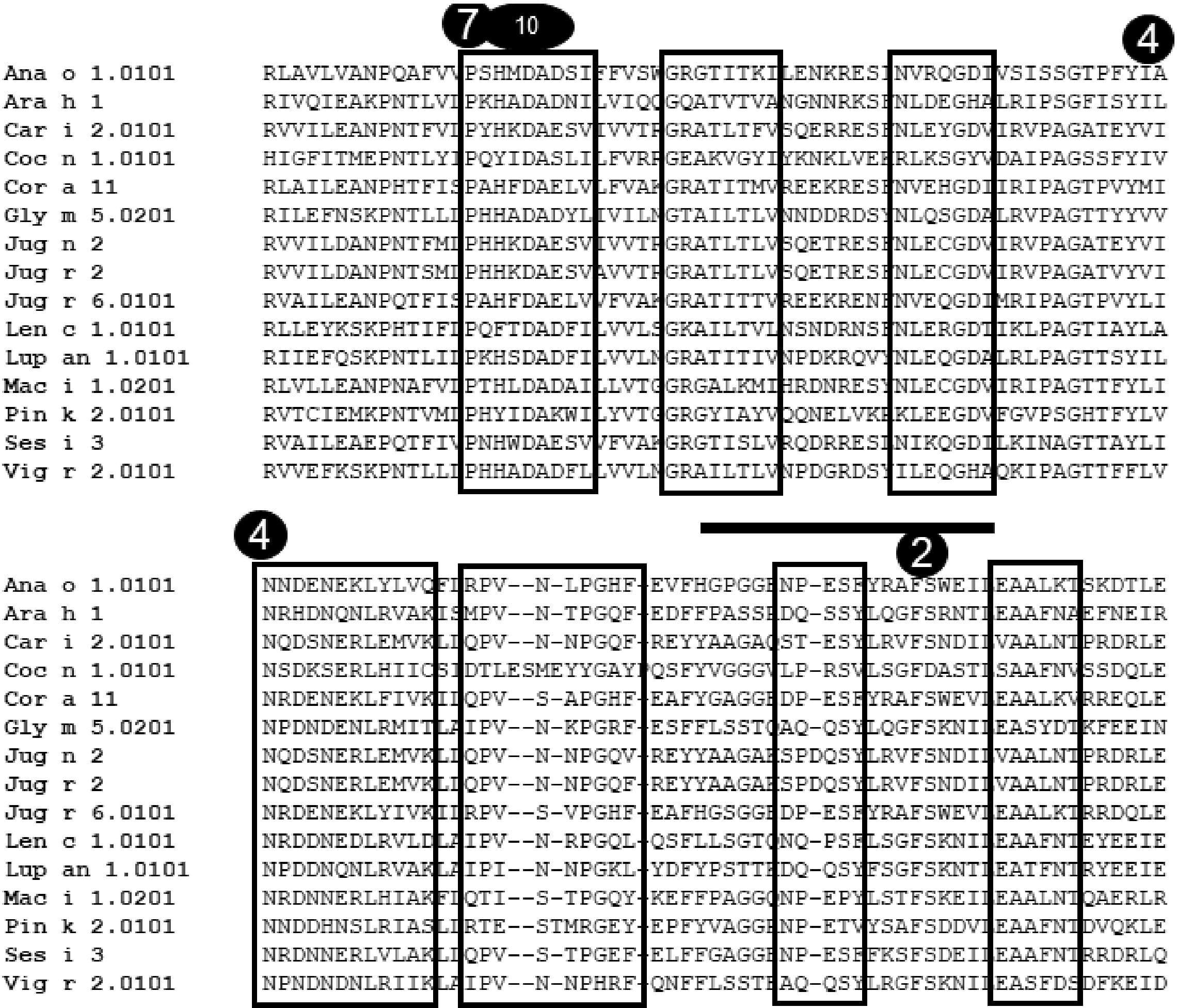
Position of conserved PCP motifs in the vicilin allergens in SDAP. (shown in black boxes). Location of local FIS peak position in Ana o 1 shown by number inside a circle, and position of Ana o 1 epitope 1 shown by a horizontal line. This figure shows the overlap of some of the FIS peak positions with the PCP motifs and epitope location, and confirms the importance of these region in the vicilin allergens.

### Finding potential novel allergens by AllergenAI

We used the Cupin protein family as annotated in the PFAM data base (PF00190) with 27,182 protein sequences as an example to show the usefulness of AllergenAI to find previously unrecognized, novel allergens that could cause adverse allergic reactions in sensitized individuals. This diverse superfamily of proteins contains the well-characterized vicilin allergens in peanut, walnut, soybean, and sesame seeds in addition to many non-allergens[27]. Some of the well-known allergens from cupin family in SDAP are shown in the Table 2. Despite very low sequence similarity, all proteins from cupin family are predicted to have a similar beta barrel fold.

We determined Allergen AI scores for a data set of cupin proteins (derived as described in Methods) that are very different from the vicilin sequences in SDAP. To find novel potential allergens, we selected the top 50 predicted entries. These sequences are from bacteria or plants. We focused on predicted sequences from plants as they may pose a higher risk of allergenicity. Based on the list of top 50 predictions, we found four potential allergens(Table 4). Three of these proteins were identified as having allergenic potential by AllerPred [17] while the AllergenFP [28] server predicted one of them. These sequences have less than 35% sequences identity with the allergens in the SDAP but have moderate E-value ( >0.1), suggesting that small modification in these proteins may affect potential allergenicity.

**Table 4:**
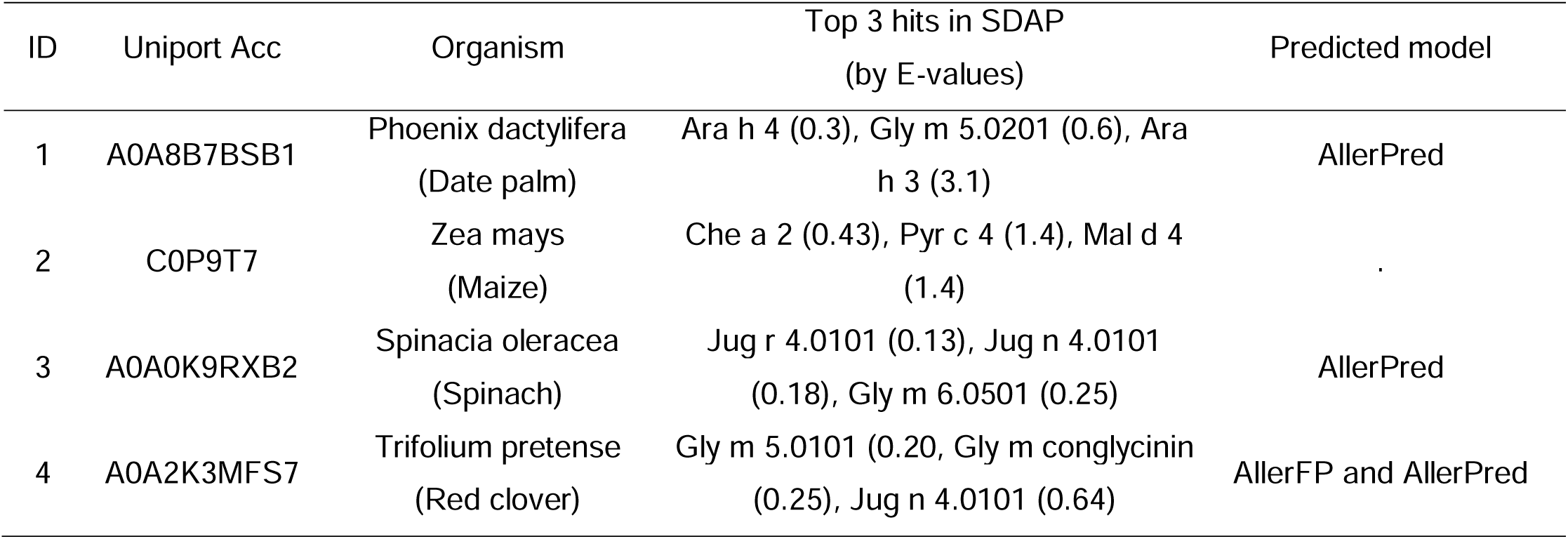
Potential allergenic proteins predicted as allergens from the Cupin family (Pfam ID PF00190) using AllergenAI.

### Protein 3D structure information enhance the performance predicting the allergenicity

We also assessed the impact of including limited 3D protein structure information for predicting allergenicity. We downloaded approximately 1500 known allergens from the SDAP for a positive training data. A negative dataset of non-allergenic proteins was generated as described in the method section. We created a one-hot encoded matrix based on the protein sequences of a balanced set, which included equal numbers (1,552) of allergenic and non-allergenic proteins. We then applied standard evaluation using five-fold cross-validation by splitting the data into a training (80%) and test data set (20%). This resulted in an allergenicity model that incorporated protein sequence information but not the 3D protein structure information. To develop an allergenicity model with both protein sequence and 3D structure information, we used the structure assignments of all proteins using the DSSP program to assign the secondary structure properties (helix, beta-strand, loop) and tertiary properties (inside/outside) of amino acid residues in individual proteins of both positive and negative data sets. We then added four additional columns to the input matrix to represent these structural properties. Using the same method of prediction quality, we found that the model with the 3D structure information (0.96/0.83 accuracy for training/testing) had a better allergenicity prediction performance than the one without 3D information (0.93/0.81) (Figure 6). Although the improvement was small, it indicated the potential benefit of utilizing the 3D protein structure information. In the future, we will use 3D structure information for all 9K positive and 9K negative proteins, to further develop AllergenAI.

**Figure 6.**
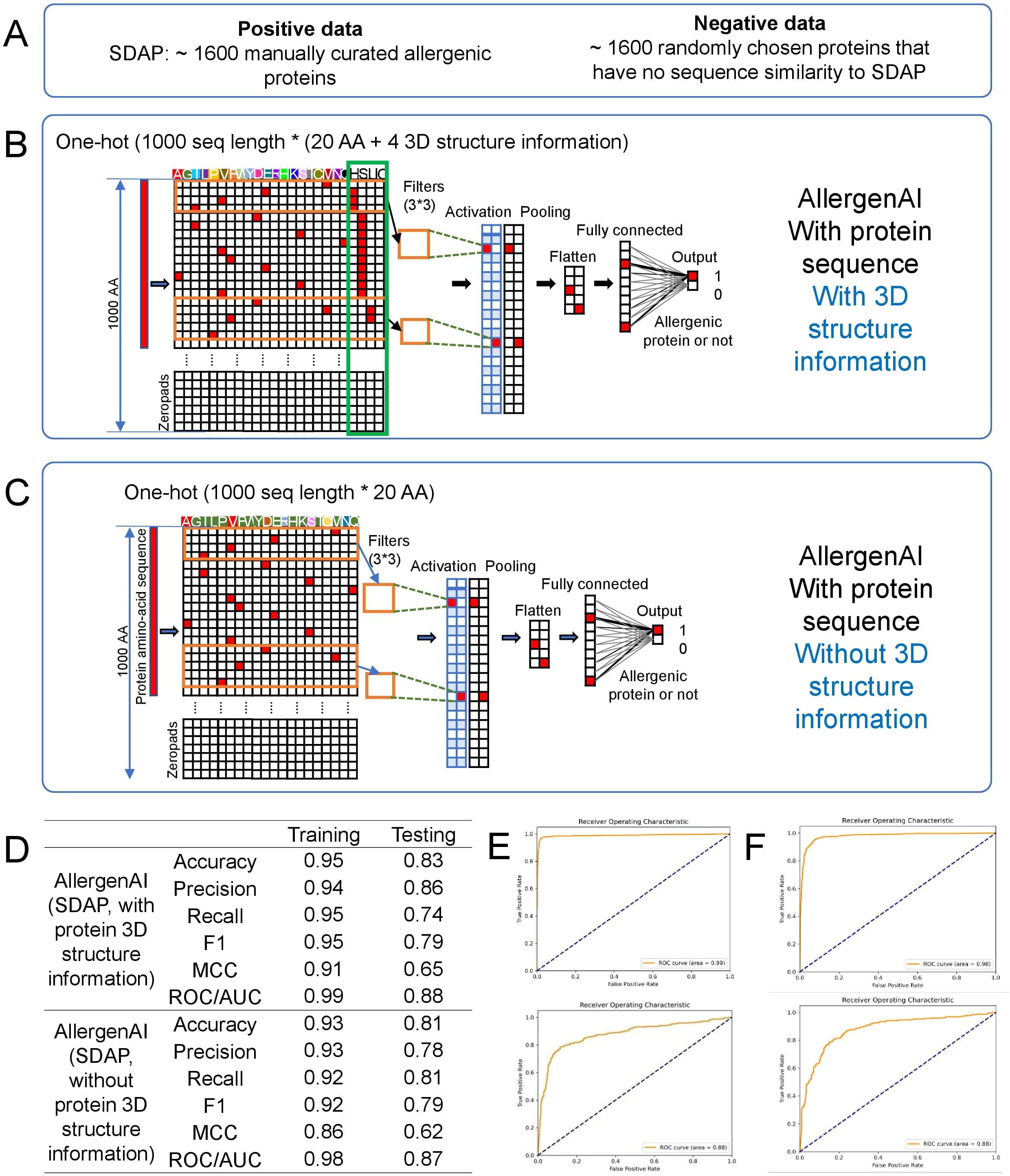
Impact of the protein 3D structure information to predict the allergenicity. (A) Training and test data composition (B) Input format and overview of the CNN model trained from the protein sequence and 3D structure information. (C) Input format and overview of the CNN model trained from the protein sequence without 3D structure information. (D) Performance comparison between these two models.

## DISCUSSION

Allergies constitute a pervasive health concern globally, with their incidence continuing to rise. They range from mild irritations to life-threatening anaphylaxis, underscoring the need for proactive measures to prevent allergic reactions. Accurately predicting allergenic proteins can contribute to designing safer food products and developing novel therapeutic strategies. Furthermore, improved prediction models can also assist in characterizing newly discovered proteins, thereby enhancing our understanding of allergenicity at a molecular level. Ultimately, advances in this domain could lead to a more informed approach to allergy management, reducing the health burden associated with allergic diseases and promoting overall patient well-being.

While allergens populate only a fraction of known protein families [14], not all proteins in the same PFAM are allergenic. Our results for the vicilins show how sequence-based ML can distinguish allergens from non-allergens in even a very large PFAM[29], and identify common features in very diverse protein sequences. We note however that even more subtle features may characterize PFAM with fewer known allergens. We also showed that AllergenAI can be used for screening large proteome databases to find potential allergens (Table 3). The four candidates with a high AllergenAI score also had local sequence similarity with known allergens in the vicilin family (Table 4).

Most of the current methods for diagnosing allergenic proteins predominantly rely on the accurate mapping of IgE epitopes, which requires detailed peptide analysis[30], or high-resolution techniques [31], such as the 3D structure of an Antigen-Antibody complex solved by X-ray crystallography. Existing allergen prediction methods are limited by their reliance on relatively small datasets that contain, at most, a few hundred proteins, with a high degree of redundancy[17]. In view of this sparse data, we developed a novel method using only long protein sequence data to predict allergenic proteins. We showed, using a feature scoring method, that there were indeed subtle commonalities in the sequences of allergens. Further, from a molecular perspective, we tested the incorporation of 3D-structure data into the training sets using elements defined by the DSSP program [32] Intriguingly, the data with structural information displayed slightly improved performance across all metrics, suggesting that the inclusion of structural data could potentially fortify the deep learning prediction. To further improve AllergenAI outcomes, we plan to generate more 3D structural information for a larger protein data set.

AllerAI is a useful tool for computationally identifying potential allergens, As shown here, the FIS scanning correlated well with PCP motifs[26] and known IgE epitopes of several allergens. Avoiding these features may also help pharmaceutical and food companies to design safer, less reactive antibody treatments and hypoallergenic protein sources[33]. Identifying potential cross reacting proteins can also help physicians to advise sensitive patients on sources of allergens to avoid.

## MATERIALS AND METHODS

### Training and test data format of AllergenAI

After gathering the sequences of allergenic proteins from SDAP[14], COMPARE[16], and AlgPred2[17], we checked the distribution of the sequence length to make a uniform input matrix for our model. The majority of the sequences below 400 AA, but to cover most of them, we set the sequence length as 1000 AA as shown in **Table 1**. For the short proteins of less than 1000 AA, we used the zero-padding approach to have the same data dimension. To feed the protein sequence information into the model, we converted the protein amino acid sequences into the one-hot encoded matrix format (1000 AA protein sequence length * 20 AA combination). In total, 13K, 2.2K, and 20K protein sequences in the training data set were from SDAP, COMPARE, and AlgPred2, respectively. Since COMPARE includes only the validated allergens, we used this dataset as the external validation dataset.

### Training and test data set for assessing the importance of 3D structures in an AI model

For the positive data set, we have repeated the performance of AllergenAI analysis using structural information of all model structures of allergenic proteins in the SDAP 2.0 database. We used the 3D structure annotation information from the predicted 3D structures mentioned above of 1,552 allergen proteins from SDAP with AA sequence lengths up to 1,000 residues (positive data set) and a balanced negative data set of 1,552 non-allergen proteins with the same length. The input of the model is a sequence of 1000 AA * 20 AA one-hot encoded amino acid matrix plus four protein 3D structure information (helix, beta-strand, loop, and inside/outside information calculated by a database of secondary structure assignments (DSSP) [34]. The output contains the probabilities corresponding to allergen and non-allergen prediction. For the negative data set, non-allergenic proteins were generated from proteins with experimental 3D structures in the Protein Data Bank [35] (March 2023 release). Using the CD-HIT program [36, 37], sequences were clustered with 30% sequence identity, resulting in ∼22.5k unique clusters. We selected one representative sequence from each cluster if the sequence length was >50 amino acids. This yielded ∼20.8k protein sequences, which were searched against SDAP 2.0 to remove all potential allergenic sequences (cutoff E-value = 0.1), resulting in 8,785 unique sequences. To obtain a balanced dataset for non-allergenic protein sequences, we randomly selected 1,552 of these sequences and considered them as our negative dataset. After dividing the data sets into 80% training and 20% testing, we applied five-fold cross-validation.

### Machine Learning Classifiers

We employed a convolutional neural network (CNN)[18, 38], serving as the classifier to predict the binary response to define a given protein as allergen or non-allergen. We reshaped the whole protein sequence data to a 3-D shape as the input before feeding it to the CNN model. Our CNN model includes the convolutional layer with 2D kernel, maxpooling layers, and fully connected layers (Figure1). We used Adam optimizer and binary cross entropy loss function in Keras for model fitting. To optimize this model, we tuned parameters such as learning rates and batch size. Additionally, we explored other techniques, such as dropout and regularization, to mitigate overfitting. We compared the AllergenAI result with SVM, a machine learning model, and AllergenFP [28], an allergenicity prediction model based on descriptor fingerprints. Our current deep neural network using the protein sequence information (without protein 3D structure information) consists of one convolutional layer, one max pooling, one flattening, and two dense layers finally to the output layer. The model involves 15,306,702 parameters including both weight matrix and bias at related layers. 13.6K out of a combined 17K protein sequences (9.6K allergens and 7.4K non-allergens) were used as training and validation sets (80% for training and 20% for validation), and the rest of 3.4K was used for an independent test.

### Five-fold cross-validation

Following the random division of allergen and non-allergen datasets, we employed a conventional 5-fold cross-validation technique to train and evaluate our model using the training set. This random partitioning process was performed five times, ensuring that each training and validation set was used without repetition. To ensure our model’s performance consistency, we calculated the mean accuracy value derived from the five individual accuracy measurements. Subsequently, these five distinct training and validation sets were employed to develop learning-based prediction models and optimize their parameters. We used the Keras Tuner Library, a systematic hyper-parameter tuning process, to optimize the performance of the CNN model. The objective of the tuning was to find the optimal learning rate for the Adam optimizer, the best rate for L2 regularization, and the ideal batch size that maximizes validation accuracy. The Keras Tuner library provides hyperparameter tuning methods like Random Search, Hyperband, and Bayesian Optimization. This study used Random Search due to its effectiveness and simplicity. Performance evaluation metrics Multiple classification performance metrics method were used to evaluate the performance of the AllergenAI model, including precision, recall, accuracy, Matthew’s correlation coefficient (MCC), as shown in Figure 2.

### Generation of a non-allergenic data set from the Cupin family

We downloaded all protein sequences from the cupin Pfam family (PF00190, 27182 sequences). Cupins belong to a highly diverse superfamily of proteins, including the vicilin subfamily, with well-characterized allergens in peanut, walnut, soybean, and sesame seeds, but also non-allergens[27]. These sequences were clustered based on 95% sequence identity to remove similar sequences, and a representative sequence from each cluster was selected for the analysis. Using this approach, we obtained 15292 unique sequences with less than 95% sequence identity. Using a FASTA search and E-value cutoff 0.1, each of the representative sequences was searched against SDAP database to find homologous proteins in SDAP. Using this approach, we assumed that all sequences having E-value less than 0.1 are allergenic while those with higher E-values are most likely non-allergenic. Based on Fasta search criteria, we obtained 2775 unique sequences which are most likely to be non-allergenic as they do not have low E-value homologs in the SDAP database.

### Feature importance score calculation

After selecting the allergenic protein candidates, users can check the distribution of the feature importance scores (FIS) of each individual allergenic protein using a group of 5 amino acids. The FIS score is calculated by masking 5 amino acids at a time by setting their values to zero and measuring the change of prediction outcome caused by the masking process. This procedure is repeated for every 5 amino acid windows along the whole protein sequence. Masking the amino acids helps to understand the contribution of these residues towards the allergenic potential of a protein, as measured in terms of FIS score. The local FIS score is thus the difference between predicted AllergenAI score of a sequence and the AllergenAI score of that protein with masked residues. Large changes in the local FIS score should indicate the importance of masked region.

## DATA AND CODE AVAILABILITY

Analyzed data and results are available as supplementary data on the website of the Briefings in Bioinformatics Journal. AllergenAI model, training and test data are available on https://fermi.utmb.edu/ and https://compbio.uth.edu/AllergenAI/.

## ACKNOWLEDGEMENTS

Funding for the study was partially supported by the NIH grant R35GM138184 to Dr. Kim, the Margaret Maccallum Gage and Tracy Davis Gage Professorship to Dr. Braun and a subcontract from the University of Colorado of NIH R01 AI165866 (PI Dr. Dreskin) to Dr. Schein. The funders had no role in study design, data collection, and analysis, decision to publish, or preparation of the manuscript.

## DECLARATION OF INTERESTS

The authors declare no competing interests.

**Supplementary Figure S1. tSNE plot of different datasets of allergenic/non-allergenic proteins.** (A) tSNE plot of SDAP allergens and similar number of non-allergens. (B) tSNE plot of combined data of SDAP and AlgPred2.

**Supplementary Figure S2. A Detailed description of the AllergenAI architectures.**

**Supplementary Figure S3. The histogram of the FIS values of individual vicilin proteins.**

## KEY POINTS

- We present a novel deep learning method, AllergenAI, using sequence data from several well-established allergen data bases to distinguish allergenic proteins from non-allergenic proteins with high accuracy.
- We also showed in a pilot study that inclusion of 3D structural information can improve the accuracy of the prediction.
- We demonstrated that a feature importance score can detect regions of allergens that coincide with results from sequence motif analysis.

