## Supplementary figures and images for "AllergenAI: a deep learning model predicting allergenicity based on protein sequence"

### Supplementary Figure S1

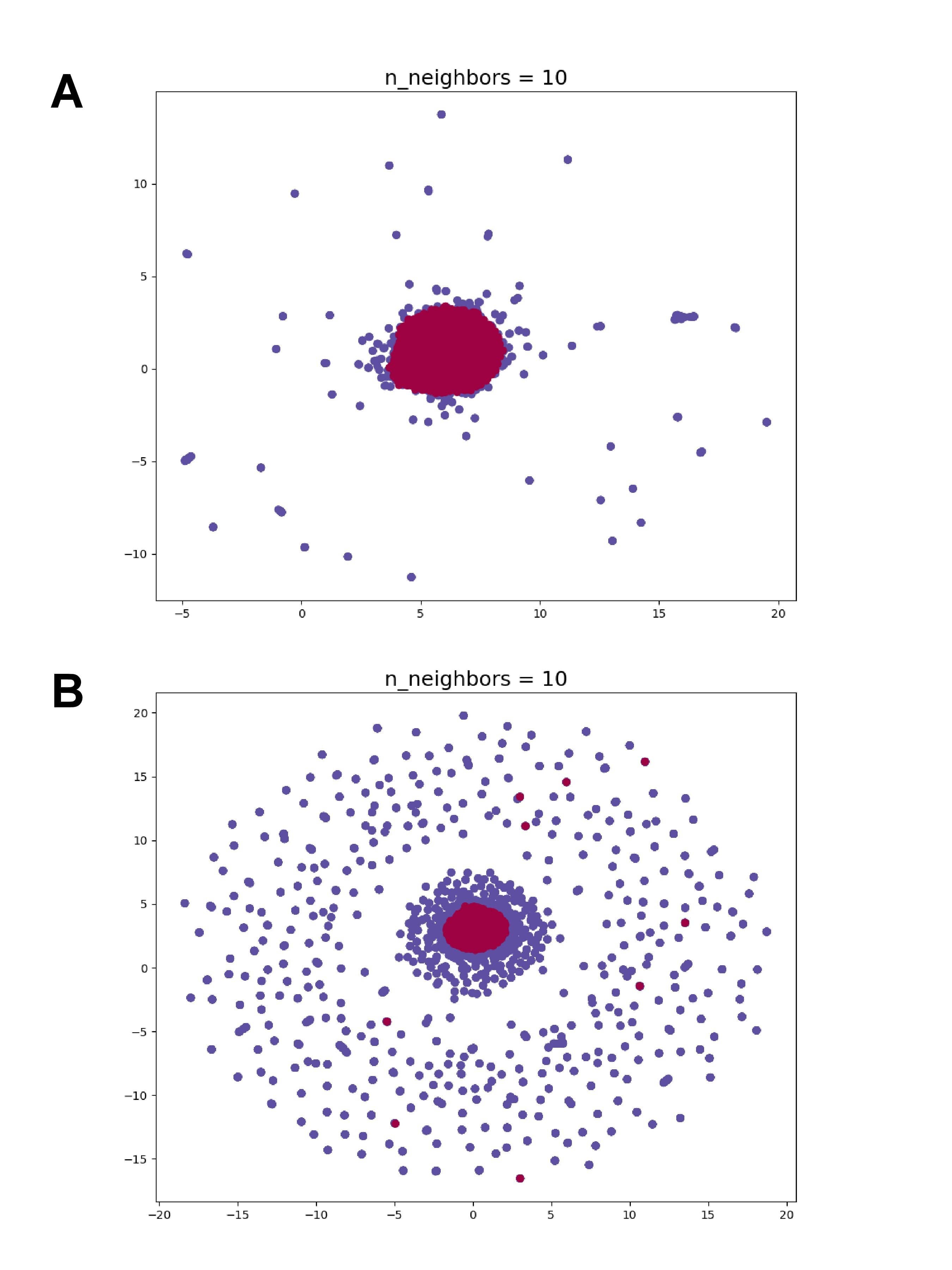

### Supplementary Figure S3

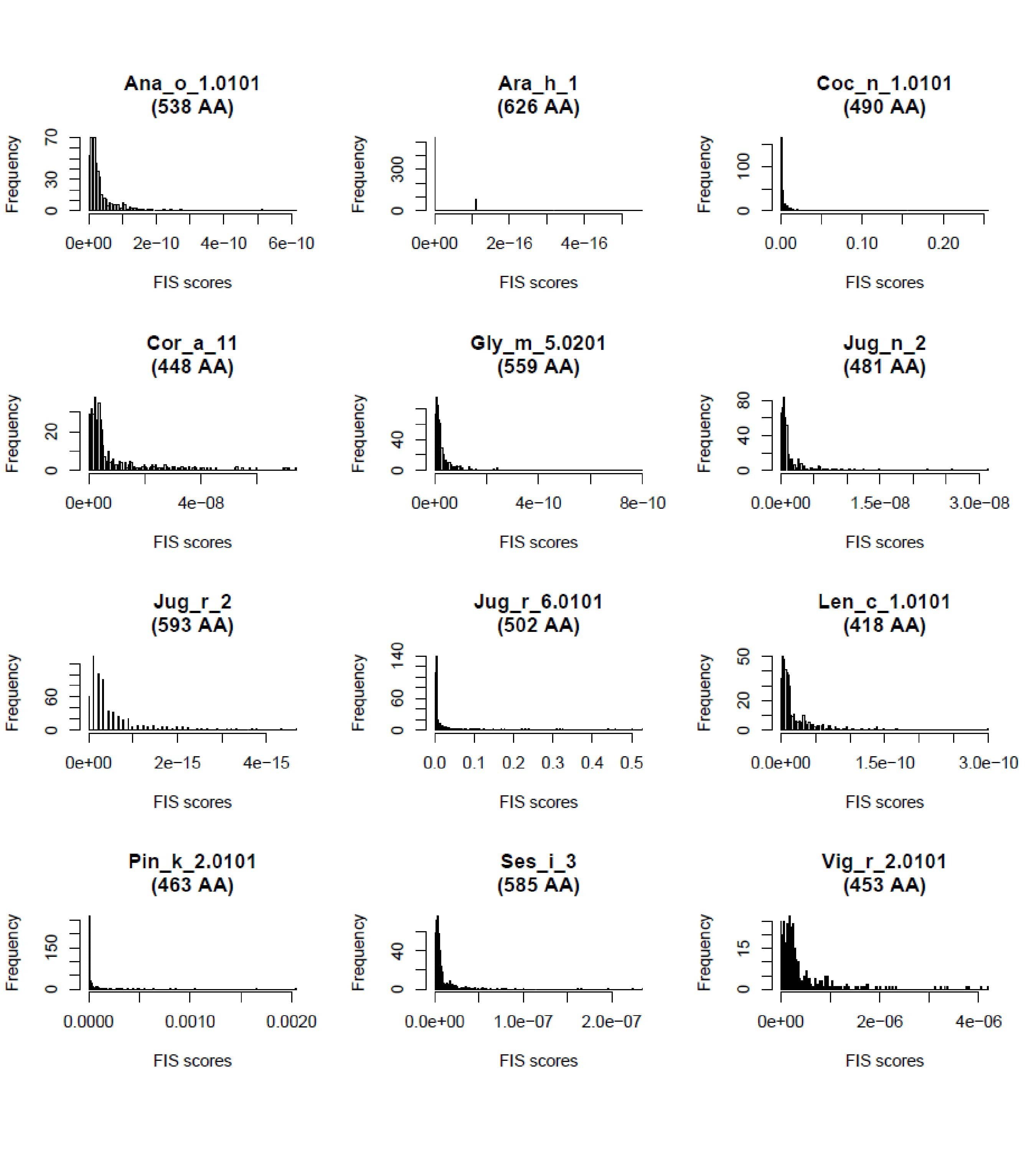

### Supplementary Figure S22

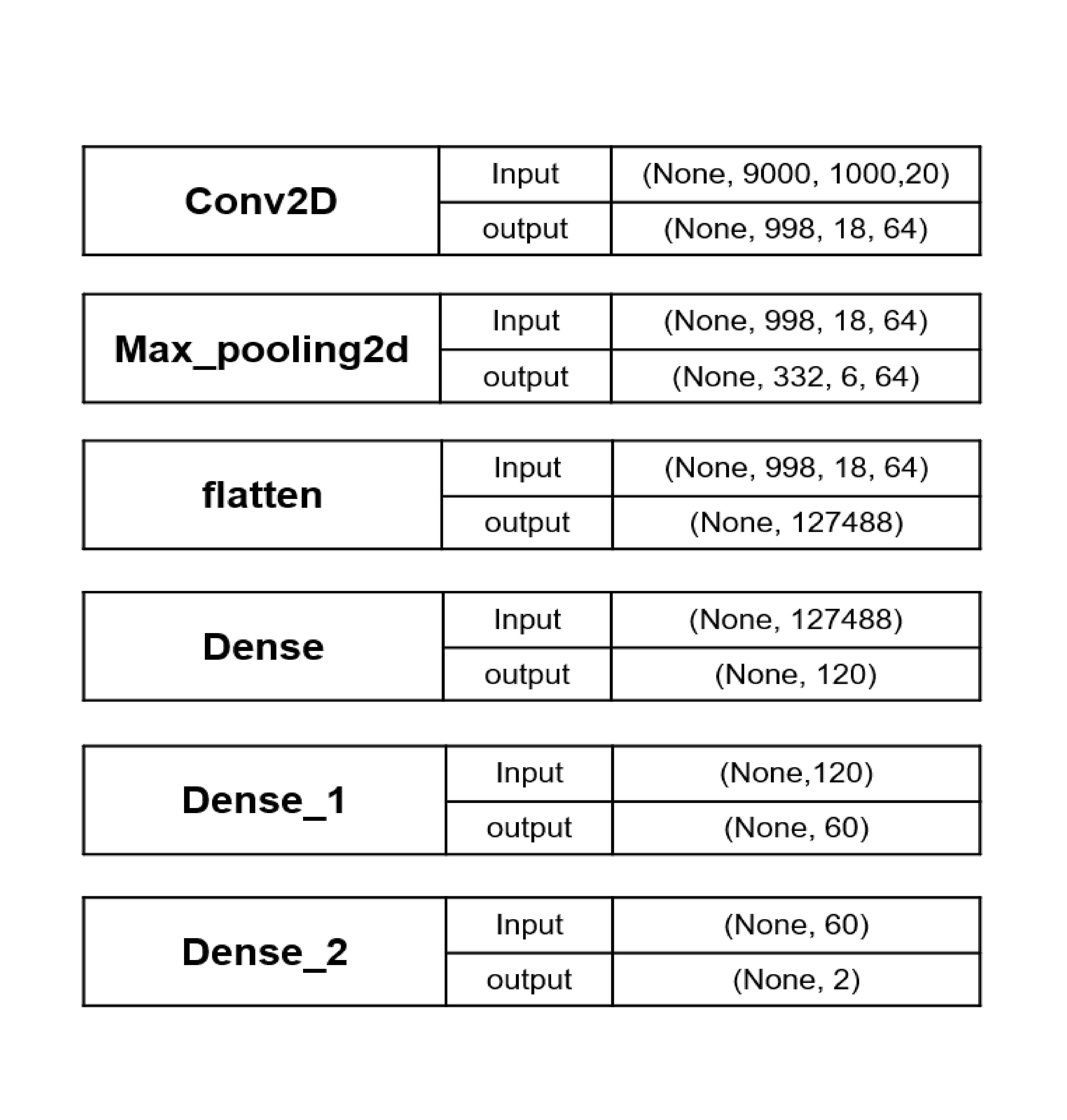
